# CD8+ T cell epitope variations suggest a potential antigen presentation deficiency for spike protein of SARS-CoV-2

**DOI:** 10.1101/2021.01.22.427863

**Authors:** Congling Qiu, Chanchan Xiao, Zhigang Wang, Xiongfei Chen, Lijuan Gao, Jieping Deng, Jun Su, Huanxing Su, Evandro Fei Fang, Zhang-Jin Zhang, Guodong Zhu, Jikai Zhang, Caojun Xie, Jun Yuan, Oscar Junhong Luo, Pengcheng Wang, Guobing Chen

## Abstract

COVID-19 is caused by a newly identified coronavirus, SARS-CoV-2, and has become a pandemic around the world. The illustration of the immune responses against SARS-CoV-2 is urgently needed for understanding the pathogenesis of the disease and its vaccine development. CD8+ T cells are critical for virus clearance and induce long lasting protection in the host. Here we identified specific HLA-A2 restricted T cell epitopes in the spike protein of SARS-CoV-2. Seven epitope peptides (n-Sp1, 2, 6, 7, 11, 13, 14) were confirmed to bind with HLA-A2 and potentially be presented by antigen presenting cells to induce host immune responses. Tetramers containing these peptides could interact with specific CD8+ T cells from convalescent COVID-19 patients, and one dominant epitope (n-Sp1) was defined. In addition, these epitopes could activate and generate epitope-specific T cells *in vitro*, and those activated T cells showed cytotoxicity to target cells. Meanwhile, all these epitopes exhibited high frequency of variations. Among them, n-Sp1 epitope variation *5L>F* significantly decreased the proportion of specific T cell activation; n-Sp1 epitope *8L>V* variant showed significantly reduced binding to HLA-A2 and decreased the proportion of n-Sp1-specific CD8+ T cell, which potentially contributes to the immune escape of SAR-CoV-2.

## Introduction

COVID-19 is an emerging pandemic sweeping the world, which is caused by SARS-CoV-2 infection (1). With no highly effective clinical treatment available for COVID-19 so far, the clearance of SARS-CoV-2 was considered relying on host immune system, especially adaptive immunity (2). It has been demonstrated that both T and B cell clones are highly expanded in the recovery phase of COVID-19 patients (3). Besides the discovery of neutralizing antibodies (4), proliferation of CD8+ T cells has been observed in the lungs of patients with mild COVID-19 (5). Although CD8+ T cell counts were dramatically reduced in severe COVID-19 cases, the revealed antigen specific T cell responses indicated an effective role of CD8+ T cells (6–8).

T cell epitopes are the basis for the initiation of T cell mediated immune responses. Thus, the identification of epitopes specific to SARS-CoV-2 and illustration of the corresponding T cell responses are of great relevance to the understanding of COVID-19 pathogenesis and vaccine development. Computational analysis has been carried out to predict potential T cell epitopes of SARS-CoV-2 (9), and T cell responses specific to SARS-CoV-2 have also been tested in convalescent COVID-19 patients by using predicted peptide ‘megapools’ (8). Recently, multiple specific CD8+ T cell epitopes have been identified throughout SARS-CoV-2 ORFs including spike protein (10). These epitopes were widely shared among SARS-CoV-2 isolates and located in regions of the virus that are not subject to mutational variation. Therefore, the above epitope identification is beneficial to the development of vaccines and medicines.

With the continuous epidemics worldwide, variations in virus strains have been found in many countries. It’s reported recently that CD8+ T cell epitope variants in SARS-CoV-2 spike protein led to persistently variable SARS-CoV-2 infection with different susceptibility and severity (11). In view of the above phenomenon, we analyzed the epitope variations of the spike protein among SARS-CoV-2 strains from different geographic regions and evaluated the epitope characteristics of these variations.

## Results

### Identification of SARS-CoV-2 T cell epitopes

To narrow down the potential candidates of SARS-CoV-2 specific antigen epitopes, we first predicted potential HLA-A2 restricted CD8+ T cell epitopes of SARS-CoV-2, SARS-CoV, MERS-CoV and Coronavirus OC43, respectively. The sequences of all the potential peptides were compared with Clustal Omega and 15 candidate peptides specific to SARS-CoV-2 were selected (Table 1 and Table S1). To validate these predicted epitopes, we first checked whether they could be presented by HLA-A2 on the antigen presenting cells (APC). T2A2 is an APC with TAP deficiency, whose peptide-MHC complex would be more stabilized if the epitopes bind suitably (Figure S1A). Comparing to the well-known CD8+ T cell epitope GIL (*GILGFVFTL*) from influenza A virus (IAV), most of the predicted SARS-CoV-2 epitopes showed reasonable HLA-A2 binding (Figure 1A&B). We further checked the direct binding of these epitopes to recombinant HLA-A2 molecules. In the UV exchange peptide-MHC assay, peptides n-Sp1, n-Sp2, n-Sp9 and n-Sp10 exhibited strong binding to HLA-A2, similar to IAV epitope GIL. However, n-Sp5 and s-Sp12 could not bind to HLA-A2, while the remaining peptides showed moderate binding capability (Figure 1C).

**Figure 1:**
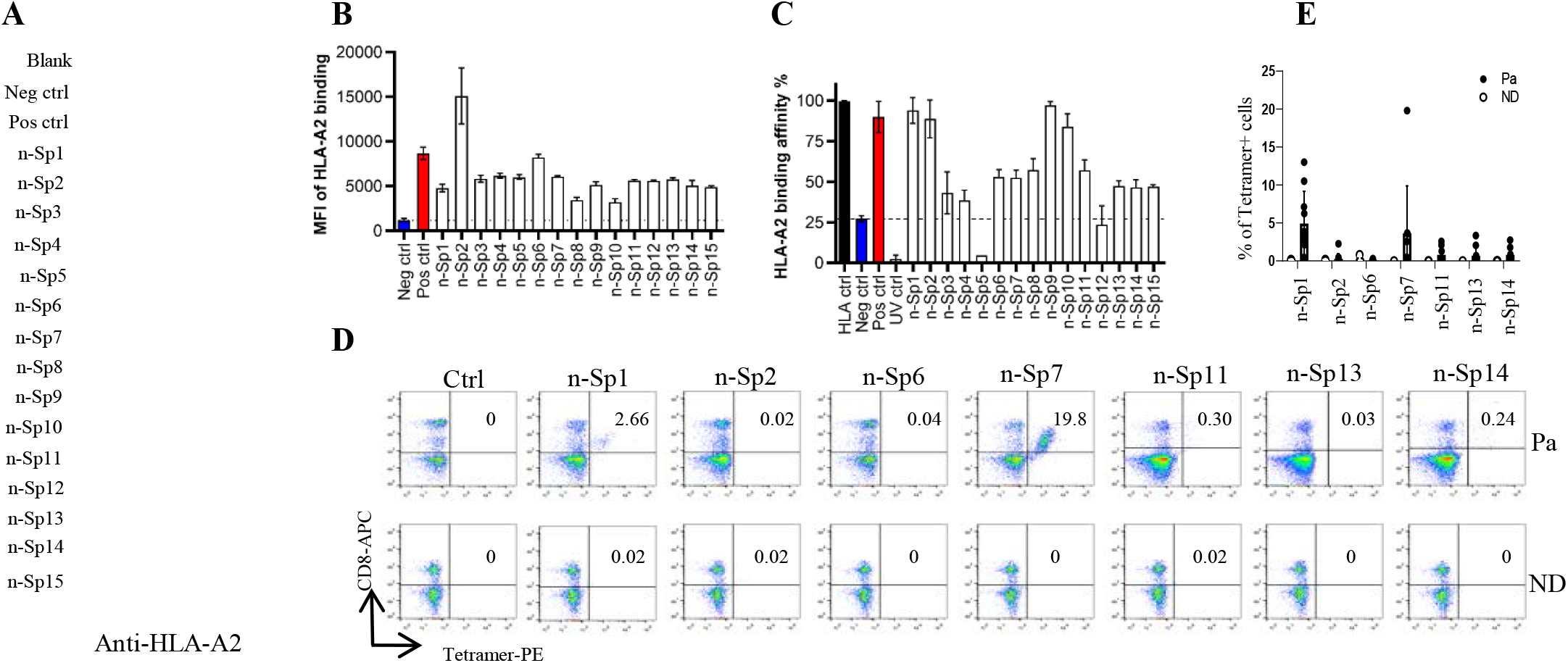
Identification of HLA-A2 restricted T cell epitopes on SARS-CoV-2 Spike protein. **A-B**: Screening of SARS-CoV-2 epitopes in T2A2 cells. Predicted peptides were synthesized and incubated with T2A2 cells. The binding of the peptide on T2A2 cells was measured with anti-HLA-A2 staining on flow cytometry. A was the representative plot of B. **C**: Screening of SARS-CoV-2 epitopes with ELISA. Peptide exchange assay was performed with coated UV-sensitive peptide/MHC complex and given peptides. The binding capability was measured with anti-HLA-A2 ELISA assay. Blank: no peptides; Neg ctrl: negative control, peptide Zika virus peptide GLQRLGYVL; Pos ctrl: positive control, influenza A M1 peptide GILGFVFTL; HLA ctrl: UV-sensitive peptide without UV irradiation; UV ctrl: UV-sensitive peptide with UV irradiation. **D-E**: Measurement of peptide specific CD8+ T cells in HLA-A2+ convalescent COVID-19 patients and healthy donors by flow cytometry. D was the representative dot plot of E. n=10. Ctrl: tetramer with UV-sensitive peptide; Pa: convalescent COVID-19 patients; ND: healthy donors.

**Table 1.** HLA-A*02:01-Restricted Peptides of S Protein Used in This Study

| Name | Start position | End position | Length | Sequence | HLA restriction | Rank | Ann_ IC50 | Ann_ rank | Smm_ IC50 | Smm_ rank | Ref |
| --- | --- | --- | --- | --- | --- | --- | --- | --- | --- | --- | --- |
| n-Sp1 | 2 | 11 | 10 | FVFLVLLPLV | A*02:01 | 0.28 | 32.64 | 0.37 | 17 | 0.2 | - |
| n-Sp2 | 133 | 141 | 9 | FQFCNDPFL | A*02:01 | 1.1 | 9.18 | 0.08 | 56.9 | 1.1 | - |
| n-Sp3 | 269 | 277 | 9 | YLQPRTFLL | A*02:01 | 0.3 | 5.36 | 0.04 | 12 | 0.3 | 1,2 |
| n-Sp4 | 386 | 395 | 10 | KLNDLCFTNV | A*02:01 | 0.42 | 15.27 | 0.14 | 50.06 | 0.7 | - |
| n-Sp5 | 417 | 425 | 9 | KIADYNYKL | A*02:01 | 0.7 | 36.12 | 0.4 | 38.82 | 0.7 | 3 |
| n-Sp6 | 612 | 620 | 9 | YQDVNCTEV | A*02:01 | 1.5 | 57.79 | 0.61 | 86.72 | 1.5 | - |
| n-Sp7 | 718 | 726 | 9 | FTISVTTEI | A*02:01 | 0.8 | 25.37 | 0.29 | 129.15 | 2 | - |
| n-Sp8 | 733 | 742 | 10 | KTSVDCTMYI | A*02:01 | 4.1 | 438.13 | 2.8 | 624.42 | 5.4 | - |
| n-Sp9 | 900 | 909 | 10 | MQMAYRFNGI | A*02:01 | 1.4 | 120.91 | 1.2 | 128.97 | 1.6 | - |
| n-Sp10 | 1000 | 1008 | 9 | RLQSLQTYV | A*02:01 | 0.7 | 16.66 | 0.17 | 36.74 | 0.7 | 3,4 |
| n-Sp11 | 1060 | 1068 | 9 | VVFLHVTYV | A*02:01 | 1.2 | 36.56 | 0.4 | 64.88 | 1.2 | 5 |
| n-Sp12 | 1095 | 1104 | 10 | FVSNGTHWFV | A*02:01 | 0.21 | 13.1 | 0.13 | 21.16 | 0.3 | - |
| n-Sp13 | 1185 | 1193 | 9 | RLNEVAKNL | A*02:01 | 0.33 | - | - | - | - | 4 |
| n-Sp14 | 1215 | 1224 | 10 | YIWLGFIAGL | A*02:01 | 0.52 | 20.48 | 0.23 | 52.42 | 0.8 | - |
| n-Sp15 | 1220 | 1228 | 9 | FIAGLIAIV | A*02:01 | 0.4 | 10.29 | 0.1 | 20.19 | 0.4 | 3,6,7 |

We then prepared pMHC tetramers with all the peptides and tested whether they could recognize antigen specific CD8+ T cells from convalescent COVID-19 patients. Overall, 31 patients were enrolled and tested (Table 2). Compared to healthy donors, the antigen specific CD8+ T cells were detected in convalescent patients with the tetramers of n-Sp1, 2, 6, 7, 11, 13 and 14, but not others. Among them, n-Sp1 and n-Sp7 based pMHC tetramers demonstrated the highest proportion of antigen specific CD8+ T cells (Figure 1D&E).

**Table 2.**
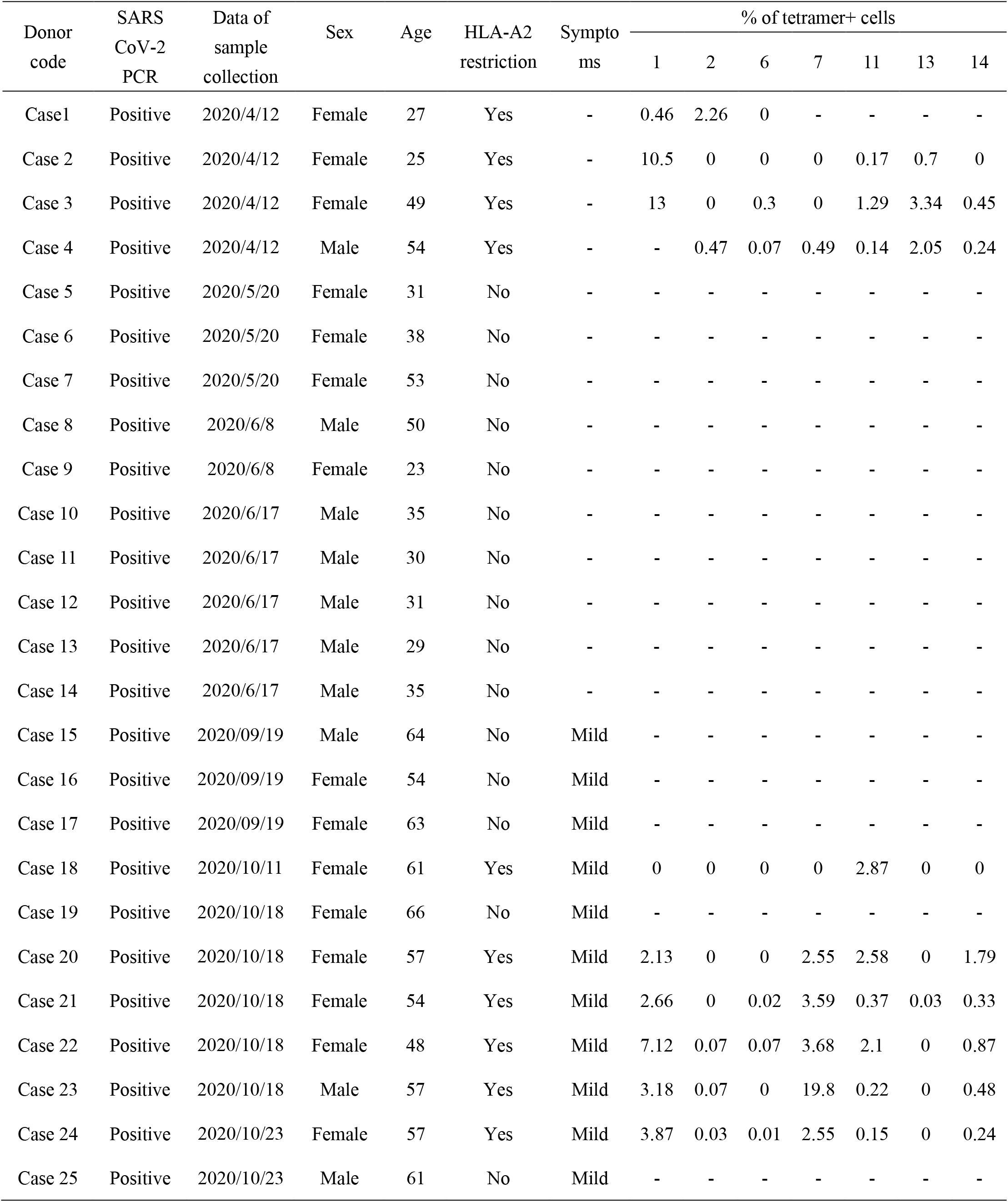
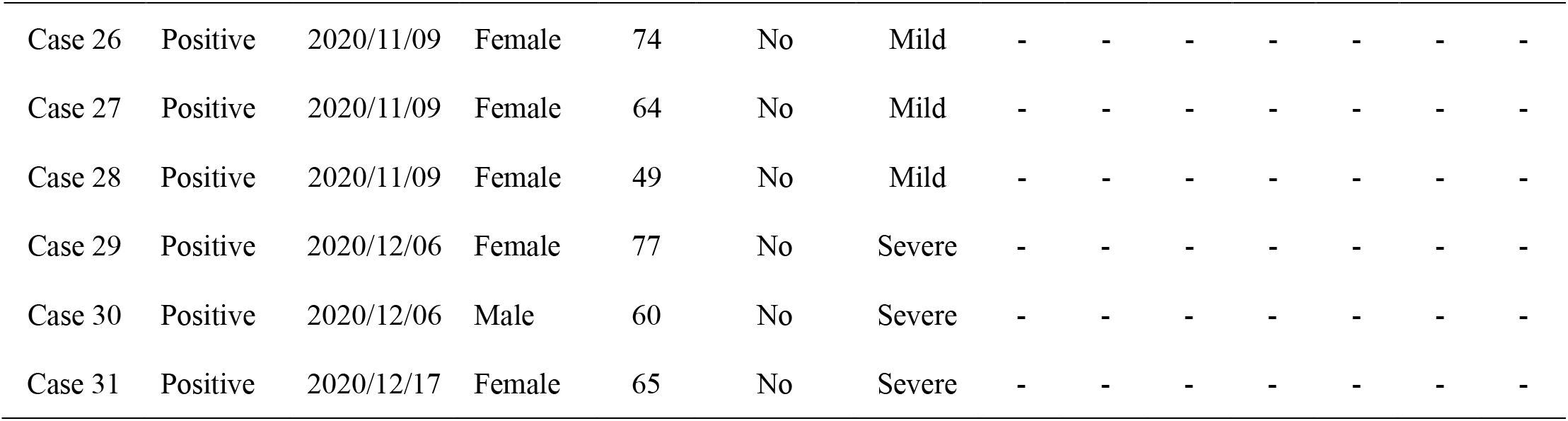
Clinical characteristics of 31 convalescent COVID-19 patients

| Donor<br>code | SARS<br>CoV-2<br>PCR | Data of<br>sample<br>collection | Sex | Age | HLA-A2<br>restriction | Sympto<br>ms | % of tetramer+ cells |  |  |  |  |  |  |
| --- | --- | --- | --- | --- | --- | --- | --- | --- | --- | --- | --- | --- | --- |
|  |  |  |  |  |  |  | 1 | 2 | 6 | 7 | 11 | 13 | 14 |
| Case1 | Positive | 2020/4/12 | Female | 27 | Yes | - | 0.46 | 2.26 | 0 | - | - | - | - |
| Case 2 | Positive | 2020/4/12 | Female | 25 | Yes | - | 10.5 | 0 | 0 | 0 | 0.17 | 0.7 | 0 |
| Case 3 | Positive | 2020/4/12 | Female | 49 | Yes | - | 13 | 0 | 0.3 | 0 | 1.29 | 3.34 | 0.45 |
| Case 4 | Positive | 2020/4/12 | Male | 54 | Yes | - | - | 0.47 | 0.07 | 0.49 | 0.14 | 2.05 | 0.24 |
| Case 5 | Positive | 2020/5/20 | Female | 31 | No | - | - | - | - | - | - | - | - |
| Case 6 | Positive | 2020/5/20 | Female | 38 | No | - | - | - | - | - | - | - | - |
| Case 7 | Positive | 2020/5/20 | Female | 53 | No | - | - | - | - | - | - | - | - |
| Case 8 | Positive | 2020/6/8 | Male | 50 | No | - | - | - | - | - | - | - | - |
| Case 9 | Positive | 2020/6/8 | Female | 23 | No | - | - | - | - | - | - | - | - |
| Case 10 | Positive | 2020/6/17 | Male | 35 | No | - | - | - | - | - | - | - | - |
| Case 11 | Positive | 2020/6/17 | Male | 30 | No | - | - | - | - | - | - | - | - |
| Case 12 | Positive | 2020/6/17 | Male | 31 | No | - | - | - | - | - | - | - | - |
| Case 13 | Positive | 2020/6/17 | Male | 29 | No | - | - | - | - | - | - | - | - |
| Case 14 | Positive | 2020/6/17 | Male | 35 | No | - | - | - | - | - | - | - | - |
| Case 15 | Positive | 2020/09/19 | Male | 64 | No | Mild | - | - | - | - | - | - | - |
| Case 16 | Positive | 2020/09/19 | Female | 54 | No | Mild | - | - | - | - | - | - | - |
| Case 17 | Positive | 2020/09/19 | Female | 63 | No | Mild | - | - | - | - | - | - | - |
| Case 18 | Positive | 2020/10/11 | Female | 61 | Yes | Mild | 0 | 0 | 0 | 0 | 2.87 | 0 | 0 |
| Case 19 | Positive | 2020/10/18 | Female | 66 | No | Mild | - | - | - | - | - | - | - |
| Case 20 | Positive | 2020/10/18 | Female | 57 | Yes | Mild | 2.13 | 0 | 0 | 2.55 | 2.58 | 0 | 1.79 |
| Case 21 | Positive | 2020/10/18 | Female | 54 | Yes | Mild | 2.66 | 0 | 0.02 | 3.59 | 0.37 | 0.03 | 0.33 |
| Case 22 | Positive | 2020/10/18 | Female | 48 | Yes | Mild | 7.12 | 0.07 | 0.07 | 3.68 | 2.1 | 0 | 0.87 |
| Case 23 | Positive | 2020/10/18 | Male | 57 | Yes | Mild | 3.18 | 0.07 | 0 | 19.8 | 0.22 | 0 | 0.48 |
| Case 24 | Positive | 2020/10/23 | Female | 57 | Yes | Mild | 3.87 | 0.03 | 0.01 | 2.55 | 0.15 | 0 | 0.24 |
| Case 25 | Positive | 2020/10/23 | Male | 61 | No | Mild | - | - | - | - | - | - | - |
| Case 26 | Positive | 2020/11/09 | Female | 74 | No | Mild | - | - | - | - | - | - | - |
| Case 27 | Positive | 2020/11/09 | Female | 64 | No | Mild | - | - | - | - | - | - | - |
| Case 28 | Positive | 2020/11/09 | Female | 49 | No | Mild | - | - | - | - | - | - | - |
| Case 29 | Positive | 2020/12/06 | Female | 77 | No | Severe | - | - | - | - | - | - | - |
| Case 30 | Positive | 2020/12/06 | Male | 60 | No | Severe | - | - | - | - | - | - | - |
| Case 31 | Positive | 2020/12/17 | Female | 65 | No | Severe | - | - | - | - | - | - | - |

To further analyze whether the epitope-bound T2A2 cells could activate T cells, we tested the expression level of T cell activation marker CD69 and the proportion of peptide-specific CD8+ T cells after stimulation with peptide-bound T2A2 cells bearing peptides n-Sp1, 2, 6, 7, 11, 13 and 14. The results demonstrated that n-Sp1, 2 and 7 had the capability to activate CD8+ T cell. In particular, n-Sp1 induced the strongest response (Figure 2). Firstly, the expression of CD69 increased significantly under the stimulation of the peptide-bound T2A2 cells (Figure 2A-B). Secondly, peptide-bound T2A2 cells stimulated T cell-mediated T2A2 killing, as the proportion of T2A2 living cells was reduced (Figure 2C-D). Thirdly, peptide-bound T2A2 cells stimulated T cell-mediated T2A2 apoptosis as the proportion of CFSE-Annexin V+ T2A2 cells was the highest (Figure 2E-F). And the expression of CD8^+^IFN-γ^+^ response was increased significantly under the stimulation of the peptide-bound T2A2 cells (Figure 2G-H). Finally, n-Sp1 based pMHC tetramers demonstrated the highest proportion of antigen specific CD8+ T cells (Figure 2I-J).

**Figure 2:**
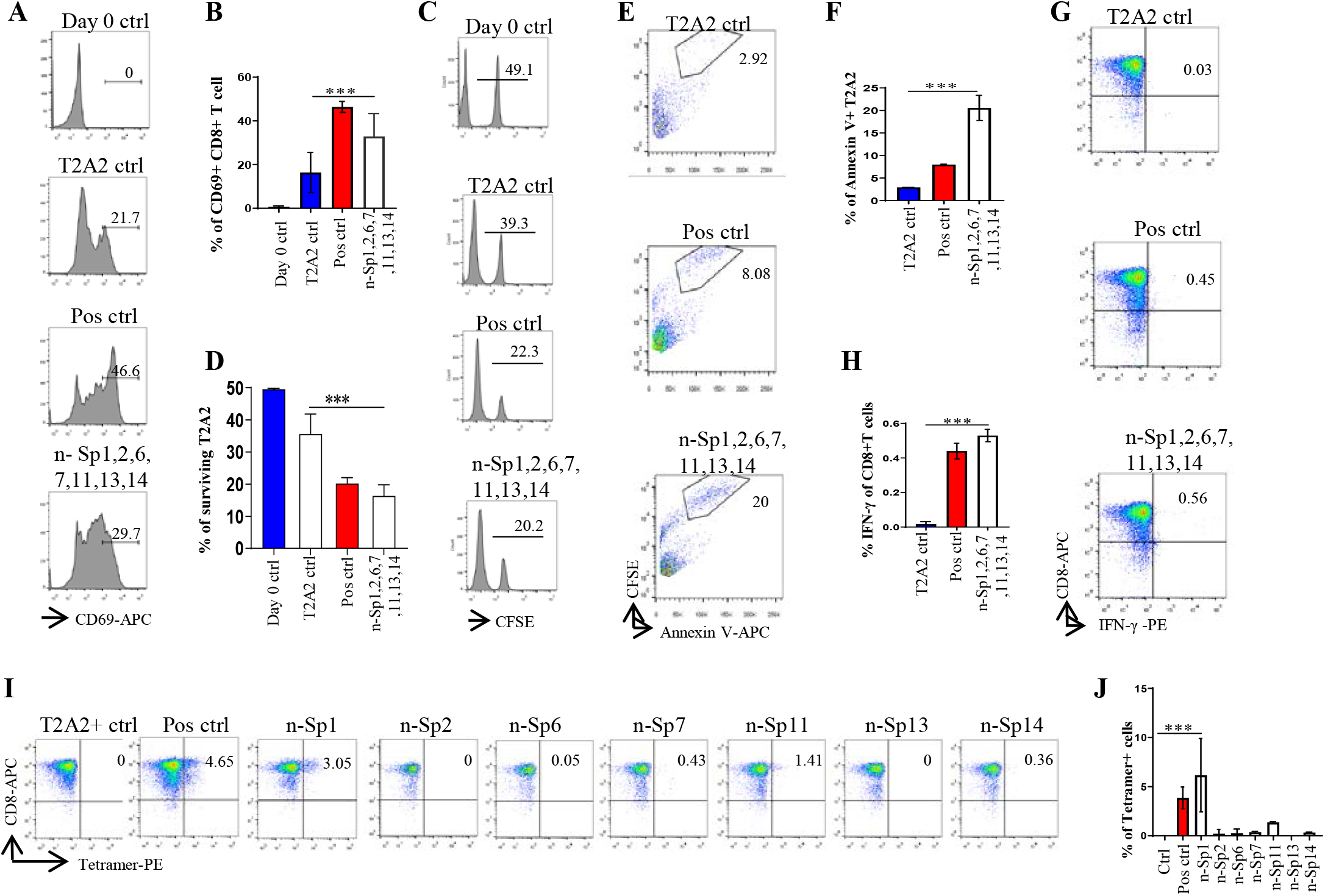
Activation of CD8+ T cell by peptides on SARS-CoV-2 Spike protein. Mitomycin pretreated T2A2 cells were loaded with n-Sp1, 2, 6, 7, 11, 13 and 14 peptides and incubated with CD8+ T cell from health donors at 1:1 ratio. Activation, cytotoxicity and generation of epitope specific CD8+ T cells were evaluated. **A-B**: Expression level of the T cell activation marker CD69 was evaluated with flow cytometry after 16 hours stimulation. A was the representative plot of B. n=3. **C-D**: Epitope specific CD8+ T cell mediated cytotoxicity was evaluated after 7 days culture. The remaining of CFSE labeled T2A2 cells was calculated as survived target cells. C was the representative plot of D. n=3. **E-F**: Apoptosis of T2A2 cells after 7 days culture. Epitope stimulated T cell mediated T2A2 apoptosis was calculated as the proportion of CFSE^+^AnnexinV^+^ cells. E was the representative plot of F. n=3. **G-H**: The expression of IFN-γ after epitope stimulation for 7 days. IFN-γ was measured with intracellular staining flow cytometry. G was the representative plot of H. n=3. **I-J**: Epitope specific CD8+ T cell generation after 7 days stimulation. The stimulated CD8+ T cells were stained with given epitope-based tetramer, and measured with flow cytometry. I was the representative plot of J. n=5. Day 0 ctrl: staining before stimulation; T2A2 ctrl: T2A2 without peptide loading; Pos ctrl: T2A2 loaded with influenza A M1 peptide GILGFVFTL.

Altogether, we therefore considered n-Sp1 to be one of the dominate CD8+ T cell epitopes specific to SARS-CoV-2.

### Analysis of Immune response in epitope variation of SARS-CoV-2

Phylogenetic analyses had indicated diverse variants of SARS-CoV-2 in global circulation (12), and therefore we examined the extent to which the above epitopes had been evolving in the 105,029 publicly available full-length SARS-CoV-2 sequences as of Sep 26^th^ 2020. We observed high frequency of sequence variation within these peptides, with n-Sp1, 2, 6, 7, 11, 13 and 14 bearing 20, 9, 13, 10, 12, 10 and 9 types of variations, respectively (Figure 3A). Among all the variations discovered, 614D>G, 613Q>H in n-Sp6 and 5L>F in n-Sp1 were the top three most frequently observed (Figure 3A).

**Figure 3:**
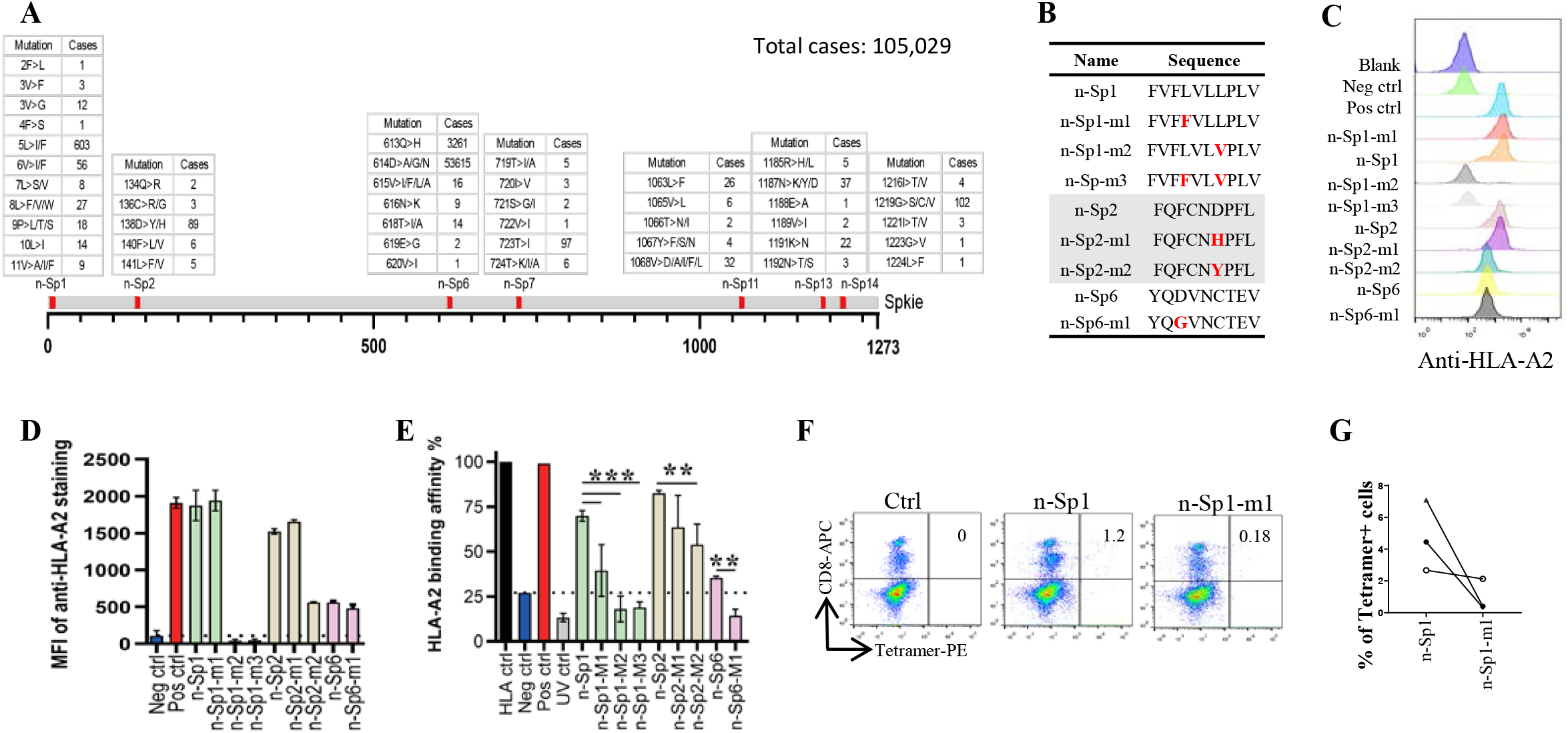
Immune response changed in epitope mutants of SARS-CoV-2. **A**: Variation frequency of epitopes on SARS-CoV-2 Spike protein. All SARS-CoV-2 virus sequences were collected from GISAID database, and the amino acid mutations in each epitope were calculated. **B**: List of the epitope mutants for following experiments. **C-D**: Comparison of epitope mutant binding to HLA-A2 in T2A2 cells. Wide type and mutated epitopes were synthesized and incubated with T2A2 cells. The peptide binding on T2A2 cells was measured with anti-HLA-A2 staining with flow cytometry. C was the representative plot of D. **E**: Comparison of epitope mutant binding to HLA-A2 with ELISA assay. Peptide exchange assay was performed with coated UV-sensitive peptide/MHC complex and given peptides. The binding capability was measured with anti-HLA-A2 ELISA assay. Blank: no peptides; HLA ctrl: UV-sensitive peptide without UV irradiation; Neg ctrl: negative control, peptide Zika virus peptide GLQRLGYVL; Pos ctrl: positive control, influenza A M1 peptide GILGFVFTL; UV ctrl: UV-sensitive peptide with UV irradiation. **F-G**: Measurement of n-Sp1-m1 specific CD8+ T cells in HLA-A2 convalescent COVID-19 patients with flow cytometry. F was the representative dot plot of G. n=3. Ctrl: tetramer with UV-sensitive peptide.

To examine how these variations might affect epitope properties, we tested the binding capability of mutated epitope peptides to HLA-A2 (Figure 3B-E). For n-Sp1, the mutant FVF*F*VLVPLV (5L>F, n-Sp1-m1) showed no change in T2A2 stabilization, but mild decrease in pMHC binding capability. The other mutant FVFLVL*<u>V</u>*PLV (8L>V, n-Sp1-m2), however, demonstrated a significant decrease in pMHC binding capability. The dual mutant of 5L>F and 8L>V (n-Sp1-m3) further confirmed the importance of L8 for the epitope properties of n-Sp1. Meanwhile, the mutants of n-Sp2 showed various levels of decreased pMHC binding capability, especially FQFCN*Y*PFL (138D>Y, n-Sp2-m2). In addition, the n-Sp6 mutant resulted in decreased pMHC binding capability, however, it showed similar T2A2 stabilization ability to the wild type peptide (Figure 3C-E).

By using tetramers containing the mutated epitopes, no CD8+ T cells specific to n-Sp1 mutation were detected in convalescent COVID-19 patients from Guangzhou, China (Figure 3F-G). However, n-Sp1 mutants was able to increase the expression of T cell activation marker CD69 (Figure 4A-B) and generate CD8+ T cells specific to the mutants (Figure 4C-D), even though the proportion of CD8+ T cells specific to mutation was less than that to wild type in the same host (Figure 4D). More importantly, although n-Sp1-m1 could activate T cells, the wild type n-Sp1 tetramer could not recognize antigen specific CD8+ T cells induced by n-Sp1 mutants (Figure 4E-F).

**Figure 4:**
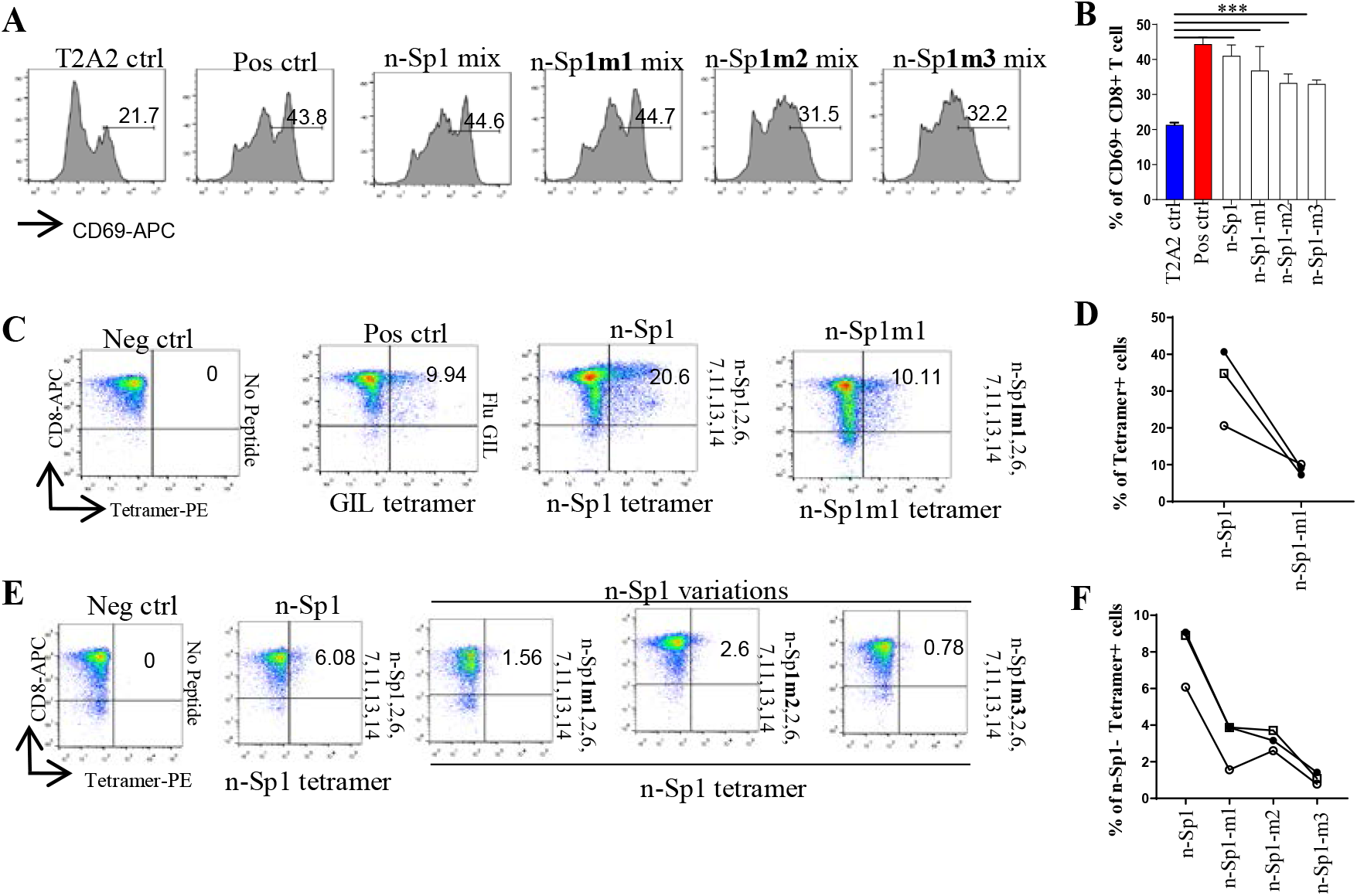
Comparison of T cell activation and function by n-Sp1 mutants. Mitomycin pretreated T2A2 cells were loaded with n-Sp2, 6, 7, 11, 13, 14, n-Sp1 and n-Sp1 mutant peptides, respectively. Next, n-Sp1 or n-Sp1 mutant peptide loaded T2A2 cells were mixed with other peptide loaded T2A2 cells and co-cultured with CD8+ T cells from health donors at 1:1 ratio. Activation, cytotoxicity and generation of n-Sp1 and its mutant specific CD8+ T cells were evaluated. **A-B**: Expression level of T cell activation marker CD69 was evaluated with flow cytometry after 16 hours stimulation. A was the representative plot of B. n=4. **C-D**: Generation of n-Sp1 and n-Sp1m1 specific CD8+ T cells. The epitope specific CD8+ T cells from the same subjects were measured with corresponding epitope peptide-based tetramer after 7 days stimulation. C was the representative plot of D. n=3. **E-F**: n-Sp1 tetramer could not recognize mutant stimulated CD8+ T cells. n-Sp1 tetramer was used to stain n-Sp1 and mutant stimulated CD8+ T cells from the same subjects after 7 days. E was the representative plot of F. n=3. Neg ctrl: T2A2 cells without peptide loading; Pos ctrl: T2A2 cells loaded with influenza A M1 peptide GILGFVFTL.

Overall, these results indicated that these emerged variations might have caused a deficiency in the antigen presentation of the dominant epitopes, which was required to rebuild a new CD8+ T cell immune response in COVID-19 patients.

## Discussion

Similar to SARS-CoV and MERS-CoV, SARS-CoV-2 has been demonstrated to induce T cell responses mainly to spike protein (13). Our study identified HLA-A2 restricted CD8+ T cell epitopes on SARS-CoV-2 spike protein. We first predicted 15 potential HLA-A2 restricted CD8+ T cell epitopes. Among them, n-Sp11, n-Sp13 and n-Sp15 have also been reported as SARS-CoV-specific epitopes and have been shown to stimulate specific CTL responses (14, 15). In the study of *Ferretti et al*., memory CD8+ T cells from convalescent COVID-19 patients could be activated by 29 epitope peptides, among which only 3 peptides were in spike protein, including n-Sp3 (10). In our study, n-Sp3 also showed a certain HLA-A2 binding capability, but it was much lower than that of the dominate epitope n-Sp1. Although their ability to activate memory CD8+ T cells from convalescent COVID-19 patients were not tested, our results did show that mixed epitope-loaded antigen presentation cells could activate T cells from healthy donors. Notably, the proportion of n-Sp1-specific T cells produced by mixed epitope activation was even higher than that of the positive control (Influenza A M1 peptide *GILGFVFTL*). Combining the results of HLA-A2 binding capability, the proportion of antigen-specific CD8+ T cells in convalescent COVID-19 patients and the ability to activate T cell, we therefore proposed n-Sp1 as the dominate CD8+ T cell epitope specific to SARS-CoV-2.

An artificial antigen presenting cell (aAPC) system was used in this study, providing a convenience protocol to validate T cell epitopes with no need of a high biosafety level laboratory. T2A2 cell line was selected for our study with following reasons. Firstly, its intrinsic deficiency in endogenous antigen presentation made it more reliable to evaluate the presentation of exogenous epitope by HLA-A2, without the interference of endogenous epitopes. Secondly, it demonstrated an excellent capability to stimulate CD8+ T cells, with the high proportion of antigen specific CD8+ T cells generated upon the primary immune challenge. Thirdly, it could also be a target cell for cytotoxic assay after labelled with given epitope. Altogether, it is an ideal model to identify T cell epitopes for novel antigen protein, especially for the low biosafety level laboratories. By using this cell model, we demonstrated that the novel epitopes from spike protein of SARS-CoV-2 had the capability to initiate CD8+ T cell response, even in the unexposed donors. These epitopes might be candidates for vaccine development in future.

Since the virus variants have been emerging geographically around the world (Figure S2 and Table S2), especially the ongoing mutation of these epitopes, it is critical to explore how the variations could affect the transmissibility and pathogenicity of the virus. Among all the variations reported so far, 614D>G and 613Q>H in n-Sp6, and 5L>F in n-Sp1 were the three most frequently observed (Figure 3A). However, n-Sp6 showed a low proportion of antigen-specific T cells in convalescent COVID-19 patients and a lack of T cell activation ability (Figure 3, Figure S3), so it’s not considered as the dominate CD8+ T cell epitope. Although 614D>G replacement has been reported as the dominant pandemic form in an epidemiological analysis (16), the G614 peptide mediated T cell activation was not significantly altered compared to D614 peptide. The possible explanation for this phenomenon is that G614 has a stronger binding ability with Angiotensin-converting Enzyme 2 (ACE2) (17) but it’s not involved in the induction of T cell immune responses. Our result was also in accordance with the report that although G614 variant showed higher viral loads in patients, it didn’t increase disease severity (18).

Although the n-Sp1 5L>F mutant could induce T cell activation, its specific T cells were undetectable in the convalescent COVID-19 patients. The possible reason was that the patients tested here were infected with wild type virus strain during the early stage of the epidemic, and the rebuild of a new CD8 + T cell immune response to the 5L>F mutant was needed. In addition, n-Sp1 8L>V mutation not only significantly reduced its binding to HLA-A2, but also showed a decreased proportion of n-Sp1-specific CD8+ T cell, which might contribute to the immune escape of SAR-CoV-2.

Taken together, our data indicated that the variation of a dominant epitope might cause the deficiency in antigen presentation, which subsequently required the rebuild of a new CD8+ T cell immune response in COVID-19 patients.

## Materials and Methods

### Human subject enrollment

The Institutional Review Board of the Affiliated Huaqiao Hospital of Jinan University approved this study. Unexposed donors were healthy individuals enrolled in Guangzhou Blood Center and confirmed with a negative report for SARS-CoV-2 RNA RT-PCR assay. These donors had no known history of any significant systemic diseases, including, but not limited to, hepatitis B or C, HIV, diabetes, kidney or liver diseases, malignant tumors, or autoimmune diseases. Convalescent donors included subjects who were hospitalized for COVID-19 or confirmed SARS-CoV-2 infection with RNA RT-PCR assay (Table 2). All subjects provided informed consent at the time of enrollment that their samples could be used for this study. Complete blood samples were collected in acid citrate dextrose tubes and stored at room temperature prior to PBMC isolation and plasma collection.

### HLA-A2 restricted T cell epitope prediction

The spike protein sequences of SARS-CoV-2 Wuhan-Hu-1 strain (NC_045512.2), SARS-CoV GD01 strain (AY278489.2), MERS-CoV Riyadh_1_2O12 strain (KF600612.1) and human coronavirus OC43 strain (KF530099.1) were used for T cell epitope prediction with the “MHC I Binding” tool (http://tools.iedb.org/mhci). The prediction method used was IEDB Recommended 2.22 (NetMHCpan EL), with MHC allele selected as HLA-A*02:01 and HLA-A*02:06, the most frequent class I HLA genotype among Chinese population (19, 20). The spike protein sequences of all four virus strains were aligned with Clustal Omega with SARS-CoV-2 as reference, and potential peptide candidates specific to SARS-CoV-2 were selected for further validation.

### Analysis of SARS-CoV-2 spike protein variations

All available SARS-CoV-2 amino acid sequences of spike protein were collected from GISAID database (GISAID.org). The sequences of each epitope were extracted for further analysis with Microsoft Office Excel 2010 and Graphic Prism (Version 7).

### Peptide screening with T2A2 cells

All peptides were synthesized in Genscript Biotech Co., Ltd (Nanjing). T2A2 cells were seeded into 96-well plates, and then incubated with peptides at 0μM, 0.625μM, 1.25μM, 2.5μM, 5μM, 10μM and 20μM at 37 °C for 4 hours. Set DMSO as blank control, Influenza A M1 peptide (GILGFVFTL) as positive control, and Zika virus peptide (GLQRLGYVL) as negative control. Cells were stained with FITC anti-human HLA-A2 antibody at 4 °C in the dark for 30 min, and acquired in flow cytometer FACS Canto (BD).

### Expression and purification HLA-A2 protein

The HLA-A2 heavy chain and β2-Microglobulin genes were cloned into the pTXBI vector through SpeI and NdeI restriction sites. Both constructs were confirmed by DNA sequencing (Shanghai Sangon Biotech, Co., Ltd). Under a heat shock at 42 °C, the constructs were transformed into *E. coli* BL21, and single colonies were picked and cultured in 2mL LB broth containing 0.1 mg/mL ampicillin at 37 °C and 370 rpm for 16 hours. The culture was subsequently expanded to 400mL and induced with 0.1mM IPTG at 24 °C for 4 hours. The cells were harvested and lysed, followed with 50 min sonication on ice. The lysate was collected into chitin resin balanced with Column Buffer (20mM Tris-HCl pH8.5, 0.5M NaCl, 1mM EDTA, 0.05%Tween20) at a low flow rate (0.5mL/min). After 15 column volume washing, on-column cleavage was conducted by quickly washing the column with 3 column volumes of the Cleavage Buffer (Column Buffer supplemented with 50 mM DTT). The column was incubated at 4 °C for 36 hours and the target protein was eluted using 3 column volumes of Column Buffer. Purified heavy chain portion was incubated with 50mM biotin (Thermo Fisher, Cat#B20656) at 4-8 °C overnight for biotinylation. Samples of every step were collected for SDS-PAGE analysis.

### Preparation of Peptide-HLA-A2 monomer

UV-sensitive peptide-HLA-A2 monomer was refolded in refolding buffer (100mM Tris-Cl pH 8.3, 5M urea, 0.4M arginine HCl, 3.7mM cysteamine, 6.3mM cysteamine, and 2mM EDTA) at 4 °C. In brief, 500μL of 20mM UV-sensitive peptide VPLRPM’J’Y (J represents Fmoc-(S)-3-amino-3-(2-nitrophenyl) propionic acid) and 30 mg of β2-Microglobulin were added to the refolding mixture via syringe. Next, 10 mg HLA-A2 heavy chain protein was added to the refolding mixture via syringe every 8 hours with 3 injections and 30 mg protein in total, and let the mixture spin in the refrigerator for 15 hours. The refolding mixture was then dialyzed against 20mM Tris (pH 8.0) for 36 hours, changing buffer about every 12 hours. Finally, the mixture was filtered and purified using weak anion exchange resin (DEAE cellulose, Santa Cruz Biotechnologies) followed by purification via size exclusion chromatography (Superdex 75pg, GE Healthcare). Samples of every step were collected for SDS-PAGE analysis.

### HLA-A2 ELISA

96-well U-bottomed plates were coated with 100μL of 0.5μg/mL streptavidin (BioLegend Cat#270302) at room temperature (18-25°C) for 16-18 hours, washed for 3 times with Washing Buffer (BioLegend Cat#421601) and blocked with Dilution Buffer (0.5M Tris pH 8.0, 1M NaCl, 1% BSA, 0.2% Tween 20) at room temperature for 30 min. Samples were prepared by 300-fold dilution in Dilution Buffer and kept on ice until usage. Set DMSO as blank control, Influenza A M1 peptide (GILGFVFTL) as positive control, and Zika virus peptide (GLQRLGYVL) as negative control. 100 μL of samples were added in duplicate and incubated for 1 hour at 37 °C. After washing three times with Washing Buffer, 100μL of diluted HRP-conjugated antibodies (BioLegend Cat#280303) were added and incubated for 1 hour at 37 °C, then washed thoroughly. 100 μL of substrate solution (10.34 mL DI Water, 1.2mL 0.1M citric acid monohydrate/tri-Sodium citrate dihydrate, pH 4.0, 240μL 40mM ABTS, 120μL hydrogen peroxide solution) was added and incubated for 8 min at room temperature in the dark on a plate shaker at 400-500 rpm. The reaction was stopped with 50μL of Stop Solution (2% w/v oxalic acid dihydrate) and absorbance at 414 nm was read using a plate reader.

### Generation of Antigen specific HLA-A2 tetramer

10mM peptide stock solution was diluted to 400μM in PBS. 20μL diluted peptide and 20μL 200μg/mL UV-sensitive peptide-HLA-A2 monomer were added into 96-well plates and mixed well by pipetting up and down. The plates were sealed, spun down at 4 °C, and exposed to UV light for 30 min on ice. The plates were then incubated for 30 min at 37 °C in the dark. 30μl of peptide-exchanged monomer and 3.3μl of PE-streptavidin (BioLegend Cat#405203) were mixed in a new plate, and incubate on ice in the dark for 30min. After the incubation, 2.4μL blocking solution (1.6μL 50mM biotin and 198.4μL PBS) to stop the reaction.

### Staining of cell-surface CD8 and tetramer

PBMC were isolated from peripheral venous blood of healthy donors and convalescent COVID-19 patients. The HLA-A2+ donors were identified by using flow cytometry. Briefly, 10^6^ PBMC were stained with FITC anti-human HLA-A2 antibody at 4 °C in the dark for 30 min, and acquired by using flow cytometer. HLA-A2 positive PBMC samples were further stained with PE labelled tetramer (home-made) and APC labelled human CD8+ antibody (BioLegend Cat#344721), and acquired with flow cytometer FACS Canto (BD).

### T cell activation

HLA-A2 expressing T2A2 cells were loaded with peptides for subsequent T cell activation. Briefly, T2A2 cells were pretreated with mitomycin for 30 min to stop cell proliferation, and incubated with given epitope peptides for 4 hours for peptide loading. 0.5×10^6^ CD8+ T cells from health donors were co-cultured with 0.5×10^6^ mixed peptide-loaded T2A2 cells stained with CFSE, and stimulated with 1μg/mL anti-human CD28 antibodies (BioLegend Cat#302901) and 50 IU/mL IL-2 (SL PHARM, Recombinant Human Interleukin-2(^125^Ala) Injection). 50 IU/mL IL-2 and 20μM peptides (n-Sp1, 2, 6, 7, 11, 13 and 14) were supplemented every two days. The T cell activation marker CD69 (BioLegend Cat#310909), tetramer specific CD8+ T cells proportion and apoptosis marker Annexin V-APC (BioLegend Cat#640919) on T2A2 cells were evaluated after 16 hours and 7 days, respectively. On day 7, cells were re-stimulated with peptides for 6 hours in the presence of GolgiPlug and GolgiStop (BD Cat#550583) plus 50 IU/mL IL-2, and the production of IFN-γ was checked with anti-IFN-γ-PE (BioLegend Cat#506507) staining.

### Statistical analysis

Graphic Prism 7 software was used for statistical analysis. One-way ANOVA was performed for group analysis. *P* values less than 0.05 were considered to be statistically significant.

## Acknowledgements

This work was supported by grants from the National Key Research and Development Program of China (2018YFC2002003), the Natural Science Foundation of China (U1801285, 81971301), Guangzhou Planned Project of Science and Technology (201904010111, 202002020039), Zhuhai Planned Project of Science and Technology (ZH22036302200067PWC) and the Initial Supporting Foundation of Jinan University. We thank Dr. Rohan Williams and Mr. Brian C. Gilmour for critical reading of the manuscript and suggestions for improvements.

## Author contributions

G.C. and P.W. designed the project. C.Q. and C.X. performed the experiments. Z.W. and X.C. analyzed the clinical information and performed sample collection. L.G., J.D., and J.Z. assisted with experiments; G.Z. J.S. C.X. and J. Y. assisted with clinical information and sample collection. C.Q., O.J.L., P.W. and G.C. analyzed the data. H.S., E.F.F. and Z.Z. assisted with data analysis. C.Q., P.W. and G.C. wrote the manuscript.

## Declaration of interests

The epitopes and tetramers from this study are the subject of a patent application.

## Reference

1. Zhu N, Zhang D, Wang W, Li X, Yang B, Song J, et al. A Novel Coronavirus from Patients with Pneumonia in China, 2019. The New England journal of medicine. 2020;382(8):727–33.

2. Zhang F, Gan R, Zhen Z, Hu X, Li X, Zhou F, et al. Adaptive immune responses to SARS-CoV-2 infection in severe versus mild individuals. Signal Transduction and Targeted Therapy. 2020;5(1):156.

3. Wen W, Su W, Tang H, Le W, Zhang X, Zheng Y, et al. Immune cell profiling of COVID-19 patients in the recovery stage by single-cell sequencing. Cell discovery. 2020;6:31.

4. Ju B, Zhang Q, Ge J, Wang R, Sun J, Ge X, et al. Human neutralizing antibodies elicited by SARS-CoV-2 infection. Nature. 2020;584(7819):15–9.

5. Liao M, Liu Y, Yuan J, Wen Y, Xu G, Zhao J, et al. Single-cell landscape of bronchoalveolar immune cells in patients with COVID-19. Nat Med. 2020;26(6):842–4.

6. Le Bert N, Tan AT, Kunasegaran K, Tham CYL, Hafezi M, Chia A, et al. SARS-CoV-2-specific T cell immunity in cases of COVID-19 and SARS, and uninfected controls. Nature. 2020.

7. Weiskopf D, Schmitz KS, Raadsen MP, Grifoni A, Okba NMA, Endeman H, et al. Phenotype and kinetics of SARS-CoV-2-specific T cells in COVID-19 patients with acute respiratory distress syndrome. Science immunology. 2020;5(48).

8. Grifoni A, Weiskopf D, Ramirez SI, Mateus J, Dan JM, Moderbacher CR, et al. Targets of T Cell Responses to SARS-CoV-2 Coronavirus in Humans with COVID-19 Disease and Unexposed Individuals. Cell. 2020.

9. Kiyotani K, Toyoshima Y, Nemoto K, and Nakamura Y. Bioinformatic prediction of potential T cell epitopes for SARS-Cov-2. J Hum Genet. 2020;65(7):569–75.

10. Ferretti AP, Kula T, Wang Y, Nguyen DMV, Weinheimer A, Dunlap GS, et al. Unbiased Screens Show CD8(+) T Cells of COVID-19 Patients Recognize Shared Epitopes in SARS-CoV-2 that Largely Reside outside the Spike Protein. Immunity. 2020;53(5):1095–107 e3.

11. Guo E, and Guo H. CD8 T cell epitope generation toward the continually mutating SARS-CoV-2 spike protein in genetically diverse human population: Implications for disease control and prevention. PLoS One. 2020;15(12):e0239566.

12. Forster P, Forster L, Renfrew C, and Forster M. Phylogenetic network analysis of SARS-CoV-2 genomes. Proceedings of the National Academy of Sciences of the United States of America. 2020;117(17):9241–3.

13. Smith TRF, Patel A, Ramos S, Elwood D, Zhu X, Yan J, et al. Immunogenicity of a DNA vaccine candidate for COVID-19. Nature Communications. 2020;11(1):2601.

14. Tsao FM, Liu HM, and Kuhl PK. Perception of native and non-native affricate-fricative contrasts: cross-language tests on adults and infants. J Acoust Soc Am. 2006;120(4):2285–94.

15. Zhou M, Xu D, Li X, Li H, Shan M, Tang J, et al. Screening and identification of severe acute respiratory syndrome-associated coronavirus-specific CTL epitopes. J Immunol. 2006;177(4):2138–45.

16. Korber B, Fischer WM, Gnanakaran S, Yoon H, Theiler J, Abfalterer W, et al. Tracking Changes in SARS-CoV-2 Spike: Evidence that D614G Increases Infectivity of the COVID-19 Virus. Cell 2020;182(4):812–27.e19.

17. Zhou B, Thao TTN, Hoffmann D, Taddeo A, Ebert N, Labroussaa F, et al. SARS-CoV-2 spike D614G variant confers enhanced replication and transmissibility. bioRxiv: the preprint server for biology. 2020:2020.10.27.357558.

18. Volz E, Hill V, McCrone JT, Price A, Jorgensen D, O’Toole Á, et al. Evaluating the Effects of SARS-CoV-2 Spike Mutation D614G on Transmissibility and Pathogenicity. Cell. 2020.

19. Gonzalez-Galarza FF, Takeshita LY, Santos EJ, Kempson F, Maia MH, da Silva AL, et al. Allele frequency net 2015 update: new features for HLA epitopes, KIR and disease and HLA adverse drug reaction associations. Nucleic Acids Res. 2015;43(Database issue):D784–8.

20. He Y, Li J, Mao W, Zhang D, Liu M, Shan X, et al. HLA common and well-documented alleles in China. HLA. 2018;92(4):199–205.

