## Supplementary material for "CD8+ T cell epitope variations suggest a potential antigen presentation deficiency for spike protein of SARS-CoV-2": Figure S1

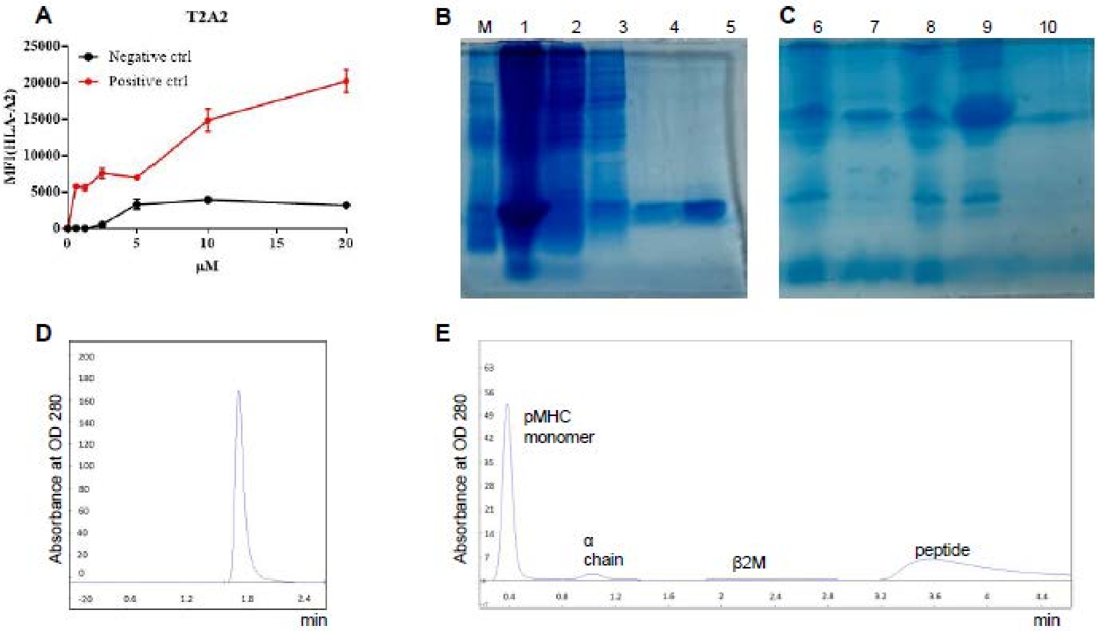


**Figure S1: Establishment of epitope validation assay.**

**A:** Establishment of T2A2 presentation assay. T2A2 cells were incubated with a series of concentrations of positive control (Influenza A M1 peptide GILGFVFTL) and negative control (Zika virus GLQRLGYVL) peptides. The stabilized HLA-A2 was detected with anti-HLA-A2 fluorescent antibody flow cytometry staining.

**B-C:** Expression and purification of recombinant protein of β2-Microglobulin (B) and HLA-A2 heavy chain (C). cDNAs of β2-Microglobulin and HLA-A2 heavy chain were subcloned into pTXB vector. The recombinant protein was expressed by IPTG induction, and then purified with chitin magnetic beads (50-70 μm paramagnetic microparticle). M represented the protein molecular weight marker; lane 1 and 6 represented the supernatant after sonication of bacteria; lane 2 and 7 represented the precipitation after sonication of bacteria; lane 3 and 8 represented flow-through fraction after chitin column loading; lane 4 and 9 represented flow-through fraction right after

Cleavage Buffer loading; lane 5 and 10 represented the elution fraction after 36 hours cleavage.

**D:** Purification of pMHC monomer complex with weak anion exchange resin (DEAE cellulose). The peak represented purified pMHC monomer.

**E:** Purification of pMHC monomer complex with size exclusion chromatography (Superdex 75pg). The four peaks represented pMHC monomer, HLA-A2 heavy chain, β2-Microglobulin (β2M) and soluble peptide, respectively.
