## Supplementary material for "CD8+ T cell epitope variations suggest a potential antigen presentation deficiency for spike protein of SARS-CoV-2": Figure S2

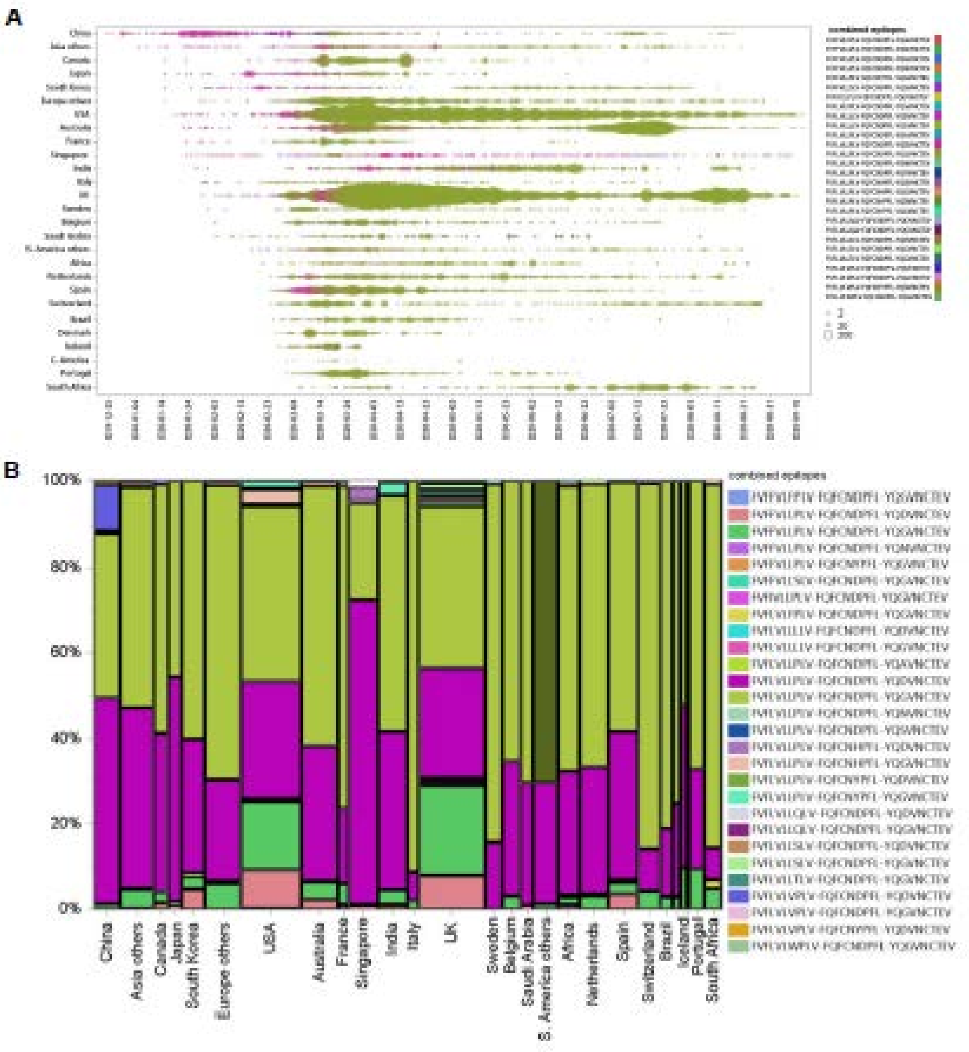


**Figure S2: The geographical features of SARS-CoV-2 dominant epitopes.**

All the sequences of n-Sp1, n-Sp2 and n-Sp6 were collected from 105,029 reported SARS-CoV-2 stains in GISAID database. The distribution of n-Sp1, n-Sp2 and n-Sp6 variants in each country with more than 500 reported cases were analyzed and presented as combination of n-Sp1, n-Sp2 and n-Sp6 characteristics. **A:** Timelines of sequence variant characteristics in various countries. **B:** The proportions of variant characteristics in various countries.
