## Supplementary material for "CD8+ T cell epitope variations suggest a potential antigen presentation deficiency for spike protein of SARS-CoV-2": Figure S3

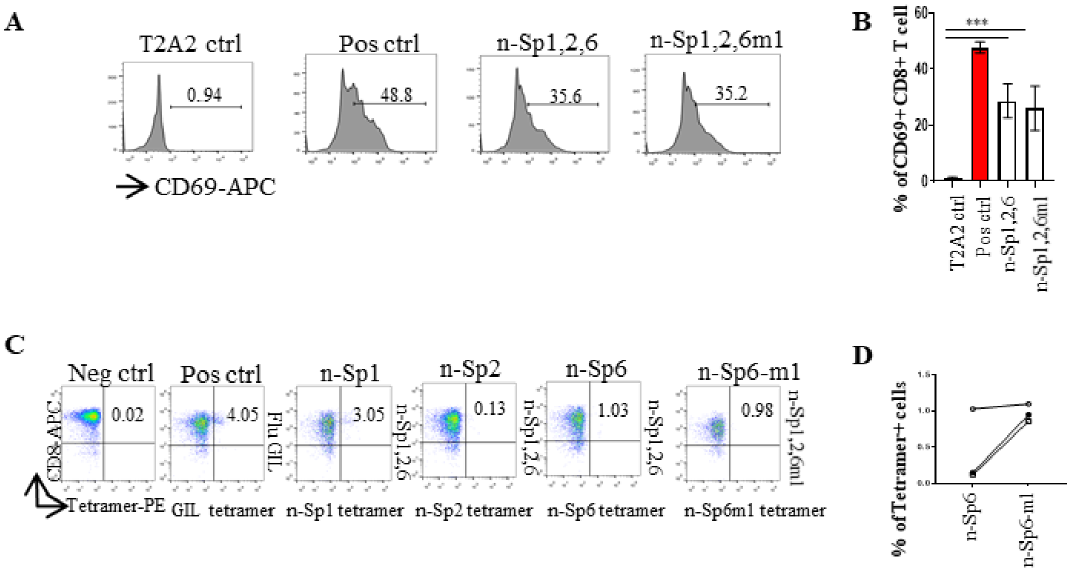


**Figure S3: Comparison of T cell activation by n-Sp6 and n-Sp6 mutant.**

Mitomycin pretreated T2A2 cells were loaded with n-Sp1, n-Sp2, n-Sp6 and peptide mutant n-Sp6m1, respectively. Next, n-Sp6 or n-Sp6m1 loaded T2A2 cells were mixed with n-Sp1/n-Sp2 loaded T2A2 cells and co-cultured with CD8+ T cells from health donors at 1:1 ratio. Epitope specific CD8+ T cell activation was evaluated.

**A-B:** Expression level of CD69 was evaluated with flow cytometry after 16 hours stimulation. A was the representative plot of B. n=3.

**C-D:** Generation of epitope specific CD8+ T cells after 7 days stimulation. The stimulated CD8+ T cells were stained with given epitope-based tetramer, and measured with flow cytometry. C was the representative plot of D. n=3.

Neg ctrl: T2A2 without peptide loading; Pos ctrl: T2A2 loaded with influenza A M1 peptide GILGFVFTL.
