## Supplementary material for "CD8+ T cell epitope variations suggest a potential antigen presentation deficiency for spike protein of SARS-CoV-2": Table S1

**Table S1: Predicted CD8+ T cell epitopes for SARS-CoV-2 virus spike protein**

| Strains | Start | End | Len | Peptide | MHC I allele | Method | Percentile | Rank | ann_ic50 | ann_rank | smm_ic50 | smm_rank | comblib_sidney2008_score | comblib_sidney2008_rank | netmhcpan_score | netmhcpan_rank |
|---|---|---|---|---|---|---|---|---|---|---|---|---|---|---|---|---|
| OC43 | 4 | 12 | 9 | ILLISLPTA | HLA-A*02:01 | Consensus | (ann/comblib_sidney2008/smm) | 1.2 | 32.02 | 0.36 | 65.63 | 1.2 | 0.0000813 | 4.7 | - | - |
| | | | | | HLA-A*02:06 | Consensus | (ann/smm) | 1.98 | 50.84 | 0.56 | 105.57 | 3.4 | - | - | - | - |
| MERS | 6 | 15 | 10 | FLLMFLLTPT | HLA-A*02:01 | Consensus | (ann/smm) | 0.2 | 18.38 | 0.2 | 6.8 | 0.2 | - | - | - | - |
| | | | | | HLA-A*02:06 | Consensus | (ann/smm) | 0.45 | 56.49 | 0.59 | 21.23 | 0.3 | - | - | - | - |
| SARS-CoV-2 | 2 | 11 | 10 | FVFLVLLPLV | HLA-A*02:01 | Consensus | (ann/smm) | 0.28 | 32.64 | 0.37 | 17 | 0.2 | - | - | - | - |
| | | | | | HLA-A*02:06 | Consensus | (ann/smm) | 0.1 | 9.2 | 0.11 | 3.67 | 0.1 | - | - | - | - |
| SARS | 2 | 10 | 9 | FIFLLFLTL | HLA-A*02:01 | Consensus | (ann/comblib_sidney2008/smm) | 0.74 | 71.52 | 0.74 | 41.03 | 0.7 | 0.000171 | 9.1 | - | - |
| | | | | | HLA-A*02:06 | Consensus | (ann/smm) | 0.81 | 100.01 | 0.93 | 20.07 | 0.7 | - | - | - | - |
| OC43 | 104 | 112 | 9 | FINGIFAKV | HLA-A*02:01 | Consensus | (ann/comblib_sidney2008/smm) | 0.9 | 20.22 | 0.22 | 49.56 | 0.9 | 0.000113 | 6.3 | - | - |
| | | | | | HLA-A*02:06 | Consensus | (ann/smm) | 0.3 | 8.84 | 0.11 | 11.44 | 0.5 | - | - | - | - |
| MERS | 110 | 118 | 9 | KQFANGFVV | HLA-A*02:01 | Consensus | (ann/comblib_sidney2008/smm) | 1.8 | 43.52 | 0.47 | 113.79 | 1.8 | 0.0000631 | 3.6 | - | - |
| | | | | | HLA-A*02:06 | Consensus | (ann/smm) | 0.19 | 7.36 | 0.08 | 5.77 | 0.3 | - | - | - | - |
| SARS-CoV-2 | | | | not found | | NA | | | | | | | | | | |
| SARS | | | | not found | | NA | | | | | | | | | | |
| OC43 | 122 | 130 | 9 | VMYSEFPAI | HLA-A*02:01 | Consensus | (ann/comblib_sidney2008/smm) | 1 | 10.83 | 0.1 | 54.21 | 1 | 0.0000586 | 3.4 | - | - |
| | | | | | HLA-A*02:06 | Consensus | (ann/smm) | 0.56 | 9.27 | 0.11 | 29.62 | 1 | - | - | - | - |
| MERS | 138 | 146 | 9 | ATIRKIYPA | HLA-A*02:01 | Consensus | (ann/comblib_sidney2008/smm) | 5.1 | 573.1 | 3.3 | 569 | 5.1 | 0.0001 | 5.7 | - | - |
| | | | | | HLA-A*02:06 | Consensus | (ann/smm) | 3.53 | 40.18 | 0.46 | 239.08 | 6.6 | - | - | - | - |
| SARS-CoV-2 | | | | not found | | NA | | | | | | | | | | |
| SARS | | | | not found | | NA | | | | | | | | | | |
| OC43 | 172 | 180 | 9 | NMCEYPHTI | HLA-A*02:01 | Consensus | (ann/comblib_sidney2008/smm) | 1.4 | 58.9 | 0.63 | 75.18 | 1.4 | 0.00231 | 53 | - | - |
| | | | | | HLA-A*02:06 | Consensus | (ann/smm) | 4.85 | 865.57 | 3.5 | 221.08 | 6.2 | - | - | - | - |
| MERS | | | | not found | | NA | | | | | | | | | | |
| SARS-CoV-2 | 133 | 141 | 9 | FQFCNDPFL | HLA-A*02:01 | Consensus | (ann/comblib_sidney2008/smm) | 1.1 | 9.18 | 0.08 | 56.9 | 1.1 | 0.0000372 | 2.1 | - | - |
| | | | | | HLA-A*02:06 | Consensus | (ann/smm) | 0.11 | 3.97 | 0.02 | 3.59 | 0.2 | - | - | - | - |
| SARS | 130 | 139 | 10 | FELCDNPFFA | HLA-A*02:01 | Consensus | (ann/smm) | 1.46 | 19.49 | 0.21 | 257.32 | 2.7 | - | - | - | - |
| | | | | | HLA-A*02:06 | Consensus | (ann/smm) | 4.65 | 260.5 | 1.8 | 428.46 | 7.5 | - | - | - | - |
| OC43 | 241 | 249 | 9 | FLFNVYLGM | HLA-A*02:01 | Consensus | (ann/comblib_sidney2008/smm) | 0.6 | 9.32 | 0.09 | 30.7 | 0.6 | 0.0000226 | 1.4 | - | - |
| | | | | | HLA-A*02:06 | Consensus | (ann/smm) | 1.59 | 80.72 | 0.79 | 71.05 | 2.4 | - | - | - | - |
| OC43 | 248 | 256 | 9 | GMALSHYYV | HLA-A*02:01 | Consensus | (ann/comblib_sidney2008/smm) | 0.5 | 11.63 | 0.11 | 24.11 | 0.5 | 0.000048 | 2.8 | - | - |
| | | | | | HLA-A*02:06 | Consensus | (ann/smm) | 1.41 | 27.87 | 0.32 | 75.43 | 2.5 | - | - | - | - |
| MERS | 277 | 286 | 10 | NMFQFATLPV | HLA-A*02:01 | Consensus | (ann/smm) | 0.2 | 17.71 | 0.19 | 15.26 | 0.2 | - | - | - | - |
| | | | | | HLA-A*02:06 | Consensus | (ann/smm) | 0.64 | 52.81 | 0.57 | 48.07 | 0.7 | - | - | - | - |

| | | | | | | | | | | | | | | | | |
|---|---|---|---|---|---|---|---|---|---|---|---|---|---|---|---|---|
| SARS-CoV-2 | | | not found | | | NA | | | | | | | | | | |
| SARS | 233 | 241 | 9 | AILTAFLPA | HLA-A*02:01 | Consensus | (ann/comblib_sidney2008/smm) | 3.5 | 168.69 | 1.5 | 284.52 | 3.5 | 0.000746 | 29 | - | - |
| | | | | | HLA-A*02:06 | Consensus | (ann/smm) | 0.76 | 9.71 | 0.12 | 41.55 | 1.4 | - | - | - | - |
| OC43 | 269 | 278 | 10 | FTLEYWVTPL | HLA-A*02:01 | Consensus | (ann/smm) | 0.61 | 29.08 | 0.32 | 60.32 | 0.9 | - | - | - | - |
| | | | | | HLA-A*02:06 | Consensus | (ann/smm) | 0.32 | 4.48 | 0.03 | 38.9 | 0.6 | - | - | - | - |
| MERS | 317 | 325 | 9 | KLQPLTFLL | HLA-A*02:01 | Consensus | (ann/comblib_sidney2008/smm) | 0.4 | 7.83 | 0.07 | 18.8 | 0.4 | 0.0000251 | 1.5 | - | - |
| | | | | | HLA-A*02:06 | Consensus | (ann/smm) | 1.27 | 20.46 | 0.25 | 70.07 | 2.3 | - | - | - | - |
| SARS-CoV-2 | 269 | 277 | 9 | YLQPRTFLL | HLA-A*02:01 | Consensus | (ann/comblib_sidney2008/smm) | 0.3 | 5.36 | 0.04 | 12 | 0.3 | 0.0000275 | 1.6 | - | - |
| | | | | | HLA-A*02:06 | Consensus | (ann/smm) | 0.96 | 16.55 | 0.22 | 50.07 | 1.7 | - | - | - | - |
| SARS | 256 | 264 | 9 | YLKPTTFML | HLA-A*02:01 | Consensus | (ann/comblib_sidney2008/smm) | 1 | 18.56 | 0.2 | 54.34 | 1 | 0.0000555 | 3.2 | - | - |
| | | | | | HLA-A*02:06 | Consensus | (ann/smm) | 2.48 | 41.92 | 0.47 | 146.07 | 4.5 | - | - | - | - |
| OC43 | 394 | 403 | 10 | KIYGMCFSSI | HLA-A*02:01 | Consensus | (ann/smm) | 1.75 | 106.19 | 1.1 | 202.53 | 2.4 | - | - | - | - |
| | | | | | HLA-A*02:06 | Consensus | (ann/smm) | 0.69 | 52.26 | 0.57 | 49.08 | 0.8 | - | - | - | - |
| MERS | 441 | 450 | 10 | LILDYFSYPL | HLA-A*02:01 | Consensus | (ann/smm) | 0.56 | 20.64 | 0.23 | 56.56 | 0.9 | - | - | - | - |
| | | | | | HLA-A*02:06 | Consensus | (ann/smm) | 0.19 | 7.64 | 0.08 | 17.66 | 0.3 | - | - | - | - |
| SARS-CoV-2 | 386 | 395 | 10 | KLNDLCFTNV | HLA-A*02:01 | Consensus | (ann/smm) | 0.42 | 15.27 | 0.14 | 50.06 | 0.7 | - | - | - | - |
| | | | | | HLA-A*02:06 | Consensus | (ann/smm) | 1.26 | 28.21 | 0.33 | 121.03 | 2.2 | - | - | - | - |
| SARS | 373 | 382 | 10 | KLNDLCFSNV | HLA-A*02:01 | Consensus | (ann/smm) | 0.62 | 14.82 | 0.14 | 80.81 | 1.1 | - | - | - | - |
| | | | | | HLA-A*02:06 | Consensus | (ann/smm) | 1.15 | 35.39 | 0.4 | 105.17 | 1.9 | - | - | - | - |
| OC43 | 425 | 433 | 9 | YLQSSNYRI | HLA-A*02:01 | Consensus | (ann/comblib_sidney2008/smm) | 0.6 | 12.4 | 0.12 | 31.49 | 0.6 | 0.0000158 | 1 | - | - |
| | | | | | HLA-A*02:06 | Consensus | (ann/smm) | 7.75 | 476.81 | 2.5 | 527.9 | 13 | - | - | - | - |
| MERS | 465 | 473 | 9 | SQFNYKQSF | HLA-A*02:01 | Consensus | (ann/comblib_sidney2008/smm) | 27 | 18607.42 | 32 | 11671.05 | 25 | 0.000658 | 27 | - | - |
| | | | | | HLA-A*02:06 | Consensus | (ann/smm) | 7 | 315.95 | 2 | 512.33 | 12 | - | - | - | - |
| SARS-CoV-2 | 417 | 425 | 9 | KIADYNYKL | HLA-A*02:01 | Consensus | (ann/comblib_sidney2008/smm) | 0.7 | 36.12 | 0.4 | 38.82 | 0.7 | 0.00016 | 8.5 | - | - |
| | | | | | HLA-A*02:06 | Consensus | (ann/smm) | 1.98 | 75.36 | 0.75 | 101.99 | 3.2 | - | - | - | - |
| SARS | 404 | 412 | 9 | VIADYNYKL | HLA-A*02:01 | Consensus | (ann/comblib_sidney2008/smm) | 0.9 | 50.27 | 0.54 | 51.65 | 0.9 | 0.000178 | 9.4 | - | - |
| | | | | | HLA-A*02:06 | Consensus | (ann/smm) | 3.35 | 152.24 | 1.3 | 183.46 | 5.4 | - | - | - | - |
| OC43 | 699 | 709 | 11 | LLFRNIKCNYV | HLA-A*02:01 | Consensus | (ann/smm) | 2.45 | 160.18 | 1.5 | 354.74 | 3.4 | - | - | - | - |
| | | | | | HLA-A*02:06 | Consensus | (ann/smm) | 3.95 | 1980.07 | 5.9 | 726.64 | 2 | - | - | - | - |
| MERS | 668 | 677 | 10 | KTHATLFGSV | HLA-A*02:01 | Consensus | (ann/smm) | 4.7 | 474.11 | 2.9 | 789.73 | 6.5 | - | - | - | - |
| | | | | | HLA-A*02:06 | Consensus | (ann/smm) | 1.01 | 62.59 | 0.63 | 78.51 | 1.4 | - | - | - | - |
| SARS-CoV-2 | 612 | 620 | 9 | YQDVNCTEV | HLA-A*02:01 | Consensus | (ann/comblib_sidney2008/smm) | 1.5 | 57.79 | 0.61 | 86.72 | 1.5 | 0.000331 | 17 | - | - |
| | | | | | HLA-A*02:06 | Consensus | (ann/smm) | 0.19 | 7.63 | 0.08 | 6.51 | 0.3 | - | - | - | - |
| SARS | 598 | 606 | 9 | YQDVNCTDV | HLA-A*02:01 | Consensus | (ann/comblib_sidney2008/smm) | 3.8 | 612.82 | 3.4 | 322.19 | 3.8 | 0.000708 | 28 | - | - |
| | | | | | HLA-A*02:06 | Consensus | (ann/smm) | 0.75 | 35.16 | 0.4 | 32.93 | 1.1 | - | - | - | - |

| | | | | | | | | | | | | | | | | |
|---|---|---|---|---|---|---|---|---|---|---|---|---|---|---|---|---|
| OC43 | 804 | 812 | 9 | FTIGNMEEF | HLA-A*02:01 | Consensus | (ann/comblib_sidney2008/smm) | 15 | 6071.63 | 15 | 3742.05 | 16 | 0.00012 | 6.6 | - | - |
| | | | | | HLA-A*02:06 | Consensus | (ann/smm) | 1.3 | 25.78 | 0.31 | 69.11 | 2.3 | - | - | - | - |
| MERS | 786 | 795 | 10 | FSFGVTQEYI | HLA-A*02:01 | Consensus | (ann/smm) | 2.25 | 401.31 | 2.6 | 153.99 | 1.9 | - | - | - | - |
| | | | | | HLA-A*02:06 | Consensus | (ann/smm) | 1.55 | 134.11 | 1.3 | 99.75 | 1.8 | - | - | - | - |
| SARS-CoV-2 | 718 | 726 | 9 | FTISVTTEI | HLA-A*02:01 | Consensus | (ann/comblib_sidney2008/smm) | 0.8 | 25.37 | 0.29 | 129.15 | 2 | 0.0000126 | 0.8 | - | - |
| | | | | | HLA-A*02:06 | Consensus | (ann/smm) | 0.6 | 8.29 | 0.1 | 32.93 | 1.1 | - | - | - | - |
| SARS | 700 | 708 | 9 | FSISITTEV | HLA-A*02:01 | Consensus | (ann/comblib_sidney2008/smm) | 0.34 | 30.13 | 0.34 | 111.97 | 1.8 | 0.00000264 | 0.2 | - | - |
| | | | | | HLA-A*02:06 | Consensus | (ann/smm) | 0.29 | 8.21 | 0.09 | 13.98 | 0.5 | - | - | - | - |
| OC43 | 820 | 828 | 9 | VTIDCAAFV | HLA-A*02:01 | Consensus | (ann/comblib_sidney2008/smm) | 0.5 | 30.83 | 0.34 | 70.81 | 1.3 | 0.00000741 | 0.5 | - | - |
| | | | | | HLA-A*02:06 | Consensus | (ann/smm) | 0.29 | 7.28 | 0.08 | 12.75 | 0.5 | - | - | - | - |
| MERS | 802 | 810 | 9 | VTVDCKQYV | HLA-A*02:01 | Consensus | (ann/comblib_sidney2008/smm) | 3.5 | 634.14 | 3.5 | 505.95 | 4.9 | 0.0000382 | 2.2 | - | - |
| | | | | | HLA-A*02:06 | Consensus | (ann/smm) | 1.87 | 73.24 | 0.74 | 91.32 | 3 | - | - | - | - |
| SARS-CoV-2 | 733 | 742 | 10 | KTSVDCTMYI | HLA-A*02:01 | Consensus | (ann/smm) | 4.1 | 438.13 | 2.8 | 624.42 | 5.4 | - | - | - | - |
| | | | | | HLA-A*02:06 | Consensus | (ann/smm) | 2.69 | 92.75 | 0.88 | 251.72 | 4.5 | - | - | - | - |
| SARS | 715 | 724 | 10 | KTSVDCNMYI | HLA-A*02:01 | Consensus | (ann/smm) | 4.9 | 864.27 | 4.2 | 646.37 | 5.6 | - | - | - | - |
| | | | | | HLA-A*02:06 | Consensus | (ann/smm) | 2.8 | 238.4 | 1.7 | 211.31 | 3.9 | - | - | - | - |
| OC43 | | | not found | | | NA | | | | | | | | | | |
| MERS | 975 | 983 | 9 | SIFYRLNGV | HLA-A*02:01 | Consensus | (ann/comblib_sidney2008/smm) | 2.5 | 52.19 | 0.56 | 174.22 | 2.5 | 0.000247 | 14 | - | - |
| | | | | | HLA-A*02:06 | Consensus | (ann/smm) | 1.02 | 18.83 | 0.24 | 51.95 | 1.8 | - | - | - | - |
| SARS-CoV-2 | 900 | 909 | 10 | MQMAYRFNGI | HLA-A*02:01 | Consensus | (ann/smm) | 1.4 | 120.91 | 1.2 | 128.97 | 1.6 | - | - | - | - |
| | | | | | HLA-A*02:06 | Consensus | (ann/smm) | 0.18 | 5.67 | 0.06 | 20.04 | 0.3 | - | - | - | - |
| SARS | 882 | 891 | 10 | MQMAYRFNGI | HLA-A*02:01 | Consensus | (ann/smm) | 1.4 | 120.91 | 1.2 | 128.97 | 1.6 | - | - | - | - |
| | | | | | HLA-A*02:06 | Consensus | (ann/smm) | 0.18 | 5.67 | 0.06 | 20.04 | 0.3 | - | - | - | - |
| OC43 | 1092 | 1100 | 9 | RLTALNAYV | HLA-A*02:01 | Consensus | (ann/comblib_sidney2008/smm) | 0.6 | 12.5 | 0.12 | 30.98 | 0.6 | 0.00000904 | 0.6 | - | - |
| | | | | | HLA-A*02:06 | Consensus | (ann/smm) | 1.01 | 28.18 | 0.33 | 47.92 | 1.7 | - | - | - | - |
| MERS | 1074 | 1082 | 9 | RLTTLNAFV | HLA-A*02:01 | Consensus | (ann/comblib_sidney2008/smm) | 0.8 | 19.74 | 0.22 | 47 | 0.8 | 0.0000313 | 1.8 | - | - |
| | | | | | HLA-A*02:06 | Consensus | (ann/smm) | 1.61 | 71.49 | 0.72 | 76.48 | 2.5 | - | - | - | - |
| SARS-CoV-2 | 1000 | 1008 | 9 | RLQSLQTYV | HLA-A*02:01 | Consensus | (ann/comblib_sidney2008/smm) | 0.7 | 16.66 | 0.17 | 36.74 | 0.7 | 0.0000109 | 0.8 | - | - |
| | | | | | HLA-A*02:06 | Consensus | (ann/smm) | 3.25 | 144.11 | 1.3 | 174 | 5.2 | - | - | - | - |
| SARS | 982 | 990 | 9 | RLQSLQTYV | HLA-A*02:01 | Consensus | (ann/comblib_sidney2008/smm) | 0.7 | 16.66 | 0.17 | 36.74 | 0.7 | 0.0000109 | 0.8 | - | - |
| | | | | | HLA-A*02:06 | Consensus | (ann/smm) | 3.25 | 144.11 | 1.3 | 174 | 5.2 | - | - | - | - |
| OC43 | 1151 | 1160 | 10 | GLYFIHFNYV | HLA-A*02:01 | Consensus | (ann/smm) | 0.17 | 13.84 | 0.13 | 12.32 | 0.2 | - | - | - | - |
| | | | | | HLA-A*02:06 | Consensus | (ann/smm) | 1.14 | 54.08 | 0.58 | 93.74 | 1.7 | - | - | - | - |
| MERS | | | not found | | | NA | | | | | | | | | | |
| SARS-CoV-2 | 1060 | 1068 | 9 | VVFLHVTYV | HLA-A*02:01 | Consensus | (ann/comblib_sidney2008/smm) | 1.2 | 36.56 | 0.4 | 64.88 | 1.2 | 0.0000617 | 3.6 | - | - |

| | | | | | | | | | | | | | | | | |
|---|---|---|---|---|---|---|---|---|---|---|---|---|---|---|---|---|
| | | | | | HLA-A*02:06 | Consensus | (ann/smm) | 0.58 | 21.97 | 0.26 | 27.14 | 0.9 | - | - | - | - |
| SARS | 1042 | 1050 | 9 | VVFLHVTYV | HLA-A*02:01 | Consensus | (ann/comblib_sidney2008/smm) | 1.2 | 36.56 | 0.4 | 64.88 | 1.2 | 0.0000617 | 3.6 | - | - |
| | | | | | HLA-A*02:06 | Consensus | (ann/smm) | 0.58 | 21.97 | 0.26 | 27.14 | 0.9 | - | - | - | - |
| OC43 | | | not found | | | | | | | | | | | | | |
| MERS | | | not found | | | | | | | | | | | | | |
| SARS-CoV-2 | 1095 | 1104 | 10 | FVSNGTHWFV | HLA-A*02:01 | Consensus | (ann/smm) | 0.21 | 13.1 | 0.13 | 21.16 | 0.3 | - | - | - | - |
| | | | | | HLA-A*02:06 | Consensus | (ann/smm) | 0.18 | 5.43 | 0.06 | 19.86 | 0.3 | - | - | - | - |
| SARS | 1077 | 1086 | 10 | FVFNGTSWFI | HLA-A*02:01 | Consensus | (ann/smm) | 0.24 | 8.67 | 0.08 | 27.7 | 0.4 | - | - | - | - |
| | | | | | HLA-A*02:06 | Consensus | (ann/smm) | 0.28 | 5.18 | 0.05 | 33.72 | 0.5 | - | - | - | - |
| OC43 | 1307 | 1315 | 9 | YVWLLICLA | HLA-A*02:01 | Consensus | (ann/comblib_sidney2008/smm) | 1.1 | 106.2 | 1.1 | 49.67 | 0.9 | 0.000527 | 23 | - | - |
| | | | | | HLA-A*02:06 | Consensus | (ann/smm) | 1.3 | 59.88 | 0.61 | 57.62 | 2 | - | - | - | - |
| MERS | 1298 | 1307 | 10 | YIWLGFIAGL | HLA-A*02:01 | Consensus | (ann/smm) | 0.52 | 20.48 | 0.23 | 52.42 | 0.8 | - | - | - | - |
| | | | | | HLA-A*02:06 | Consensus | (ann/smm) | 0.89 | 52.25 | 0.57 | 70.62 | 1.2 | - | - | - | - |
| SARS-CoV-2 | 1215 | 1224 | 10 | YIWLGFIAGL | HLA-A*02:01 | Consensus | (ann/smm) | 0.52 | 20.48 | 0.23 | 52.42 | 0.8 | - | - | - | - |
| | | | | | HLA-A*02:06 | Consensus | (ann/smm) | 0.89 | 52.25 | 0.57 | 70.62 | 1.2 | - | - | - | - |
| SARS | 1197 | 1206 | 10 | YVWLGFIAGL | HLA-A*02:01 | Consensus | (ann/smm) | 0.75 | 36.85 | 0.4 | 78.97 | 1.1 | - | - | - | - |
| | | | | | HLA-A*02:06 | Consensus | (ann/smm) | 0.82 | 30.67 | 0.35 | 74.8 | 1.3 | - | - | - | - |
| OC43 | 1313 | 1321 | 9 | CLAGVAMLV | HLA-A*02:01 | Consensus | (ann/comblib_sidney2008/smm) | 0.5 | 36.34 | 0.4 | 29.59 | 0.5 | 0.0000208 | 1.3 | - | - |
| | | | | | HLA-A*02:06 | Consensus | (ann/smm) | 1.05 | 57.79 | 0.6 | 44.42 | 1.5 | - | - | - | - |
| MERS | 1303 | 1311 | 9 | FIAGLVALA | HLA-A*02:01 | Consensus | (ann/comblib_sidney2008/smm) | 1.2 | 24.38 | 0.28 | 61.82 | 1.2 | 0.000152 | 8.2 | - | - |
| | | | | | HLA-A*02:06 | Consensus | (ann/smm) | 0.95 | 26.18 | 0.31 | 46.51 | 1.6 | - | - | - | - |
| SARS-CoV-2 | 1220 | 1228 | 9 | FIAGLIAIV | HLA-A*02:01 | Consensus | (ann/comblib_sidney2008/smm) | 0.4 | 10.29 | 0.1 | 20.19 | 0.4 | 0.0000289 | 1.7 | - | - |
| | | | | | HLA-A*02:06 | Consensus | (ann/smm) | 0.32 | 11.13 | 0.14 | 12.46 | 0.5 | - | - | - | - |
| SARS | 1202 | 1210 | 9 | FIAGLIAIV | HLA-A*02:01 | Consensus | (ann/comblib_sidney2008/smm) | 0.4 | 10.29 | 0.1 | 20.19 | 0.4 | 0.0000289 | 1.7 | - | - |
| | | | | | HLA-A*02:06 | Consensus | (ann/smm) | 0.32 | 11.13 | 0.14 | 12.46 | 0.5 | - | - | - | - |
