## Supplementary material for "CD8+ T cell epitope variations suggest a potential antigen presentation deficiency for spike protein of SARS-CoV-2": Table S2

### Table S2: The first reported mutation of SARS-CoV-2 main epitopes

| n-Sp1 | n-Sp2 | n-Sp6 | First reported date | Country |
|---|---|---|---|---|
| FVFLVLLPLV | FQFCNDPFL | YQDVNCTEV | 2019/12/24 | China |
| FVFLVLLPLV | FQFCNDPFL | YQ**G**VNCTEV | 2020/1/4 | Thailand |
| FVFLVL**V**PLV | FQFCNDPFL | YQDVNCTEV | 2020/1/30 | China-HK |
| FVFLVL**V**PLV | FQFCN**Y**PFL | YQDVNCTEV | 2020/2/9 | China-HK |
| FVF**F**VLLPLV | FQFCNDPFL | YQDVNCTEV | 2020/2/10 | Singapore |
| FVF**F**VLLPLV | FQFCNDPFL | YQ**G**VNCTEV | 2020/2/29 | UK |
| FVFLVLL**S**LV | FQFCNDPFL | YQDVNCTEV | 2020/3/2 | China |
| FVFLVLL**L**LV | FQFCNDPFL | YQDVNCTEV | 2020/3/10 | USA |
| FVFLVLLPLV | FQFCN**H**PFL | YQDVNCTEV | 2020/3/18 | UK |
| FVFLVLLPLV | FQFCN**H**PFL | YQ**G**VNCTEV | 2020/3/18 | Russia |
| FVFLVL**W**PLV | FQFCNDPFL | YQ**G**VNCTEV | 2020/3/18 | France |
| FVFLVLL**S**LV | FQFCNDPFL | YQ**G**VNCTEV | 2020/3/24 | UK |
| FVFLVLL**L**LV | FQFCNDPFL | YQ**G**VNCTEV | 2020/3/25 | Spain |
| FVFLVLLPLV | FQFCNDPFL | YQ**N**VNCTEV | 2020/4/1 | UK |
| FVF**F**VLLPLV | FQFCNDPFL | YQ**N**VNCTEV | 2020/4/2 | UK |
| FVFLVLLPLV | FQFCN**Y**PFL | YQ**G**VNCTEV | 2020/4/2 | UK |
| FVFLVLL**T**LV | FQFCNDPFL | YQ**G**VNCTEV | 2020/4/4 | UK |
| FVFLVLL**Q**LV | FQFCNDPFL | YQDVNCTEV | 2020/4/9 | India |
| FVF**I**VLLPLV | FQFCNDPFL | YQ**G**VNCTEV | 2020/4/28 | USA |
| FVFLVL**V**PLV | FQFCNDPFL | YQ**G**VNCTEV | 2020/4/29 | UK |
| FVFLVLLPLV | FQFCN**Y**PFL | YQDVNCTEV | 2020/5/3 | UK |
| FVFLVLLPLV | FQFCNDPFL | YQ**S**VNCTEV | 2020/5/21 | USA |
| FVF**F**VLL**S**LV | FQFCNDPFL | YQ**G**VNCTEV | 2020/6/24 | UK |
| FVF**F**VL**F**PLV | FQFCNDPFL | YQ**G**VNCTEV | 2020/6/26 | USA |
| FVFLVLLPLV | FQFCNDPFL | YQ**A**VNCTEV | 2020/7/13 | South Korea |
| FVF**F**VLLPLV | FQFCN**Y**PFL | YQ**G**VNCTEV | 2020/7/18 | Bangladesh |
| FVFLVLL**Q**LV | FQFCNDPFL | YQ**G**VNCTEV | 2020/7/20 | Switzerland |
